# Interchangeability of Periplasmic Adaptor Proteins AcrA and AcrE in forming functional efflux pumps with AcrD in *Salmonella* Typhimurium

**DOI:** 10.1101/2021.03.24.436855

**Authors:** Ilyas Alav, Vassiliy N. Bavro, Jessica M. A. Blair

**Affiliations:** Institute of Microbiology and Infection, University of Birmingham, Edgbaston, Birmingham, B15 2TT, United Kingdom; School of Life Sciences, University of Essex, Colchester, CO4 3SQ, United Kingdom

## Abstract

**Background:** RND efflux pumps are important mediators of antibiotic resistance. RND pumps including the principal multidrug-efflux pump AcrAB-TolC in *Salmonella*, are tripartite systems, with an inner membrane RND-transporter, a periplasmic adaptor protein (PAP) and an outer membrane factor (OMF). We previously identified the residues required for binding between the PAP AcrA and the RND-transporter AcrB and have demonstrated that PAPs can function with non-cognate transporters. AcrE and AcrD/AcrF are homologues of AcrA and AcrB, respectively. Here, we show that AcrE can interact with AcrD, which does not possess its own PAP, and establish that the residues previously identified in AcrB-binding are also involved in AcrD-binding.

**Methods:** The *acrD* and *acrE* genes were expressed into a strain lacking *acrABDEF* (Δ3RND). PAP residues involved in promiscuous interactions were predicted based on previously defined PAP-RND interactions and corresponding mutations generated in *acrA* and *acrE*. Antimicrobial susceptibility of the mutant strains was determined.

**Results:** Co-expression of *acrD* and *acrE* significantly decreased susceptibility of the Δ3RND strain to AcrD substrates showing that AcrE can form a functional complex with AcrD. The substrate profile of *Salmonella* AcrD differed from that of *E. coli* AcrD. Mutations targeting the previously defined PAP-RND interaction sites in AcrA/AcrE impaired efflux of AcrD-dependent substrates.

**Conclusions:** These data indicate that AcrE forms an efflux-competent pump with AcrD and thus presents an alternative PAP for this pump. Mutagenesis of the conserved RND-binding sites validates the interchangeability of AcrA and AcrE, highlighting them as potential drug targets for efflux inhibition.

## Introduction

Multidrug-resistance (MDR) efflux pumps play a major role in antibiotic resistance of bacteria by reducing the intracellular concentration of drugs^1, 2^. In particular, the Resistance-Nodulation-Division (RND) family of efflux pumps contribute to clinically relevant antibiotic resistance in Gram-negative bacteria, such as *Salmonella enterica*^3–6^. Tripartite RND pumps span the double membrane of Gram-negative bacteria and consist of an inner membrane RND-transporter, a periplasmic adaptor protein (PAP) and an outer membrane factor (OMF)^7–9^. Owing to their unique organisation, tripartite efflux pumps can directly export drugs across the outer membrane to the extracellular environment. The majority of RND pumps exhibit a broad substrate profile, which includes multiple classes of antibiotics, biocides, detergents, dyes and metals^10, 11^. Furthermore, there is increasing evidence that RND pumps also play a role in various biological processes and behaviours, including biofilm formation, detoxification, quorum sensing and virulence^12–14^.

In *S. enterica*, five RND pumps have been characterised: AcrAB, AcrD, AcrEF, MdtABC and MdsABC^5^. The AcrAB pump is constitutively expressed in *S. enterica* and displays a remarkably wide substrate profile, consisting of multiple classes of antibiotics, bile salts, detergents and dyes^5^. The AcrEF system possesses a similar substrate profile to AcrAB but is not constitutively expressed^5, 15^. In *S. enterica*, AcrB is 80% identical to AcrF, whereas AcrD is 64% and 65% identical to AcrB and AcrF, respectively^16, 17^. This sequence divergence is reflected in the substrate profile of AcrD, which exhibits markedly narrower substrate range compared to AcrB and AcrF. In *Escherichia coli*, AcrD has been shown to export aminoglycosides and anionic β-lactams^18–20^. Compared to AcrB, AcrD has a stronger preference for anionic β-lactams, which is linked to differences in the access binding pocket^21^. Homology modelling of *E. coli* AcrD combined with molecular dynamic simulations have also suggested that the different substrate specificities between AcrB and AcrD stem from the corresponding differences in the physicochemical and topological properties of their binding pockets^22^. Until now, this view of AcrD substrate selectivity has been assumed to also apply to the AcrD pump in *S. enterica.*

In *Salmonella,* most of the RND pumps require the OMF TolC to form a functional tripartite complex, the exception being MdsABC, which can function with either MdsC or TolC^23^. The RND-transporter genes are usually co-located with their cognate PAP on a single operon. In *S. enterica* there are four RND-associated PAPs: AcrA, AcrE, MdtA and MdsA^5^. Based on sequence analyses and structural alignments, AcrA and AcrE have been shown to be the most closely related, with a sequence identity of 69.3% over their first 374 residues as calculated by Expasy SIM server^24^, and just a single gap in the alignment of this region, which maps to the signal sequence and does not impact the mature protein. Correspondingly, AcrA and AcrE share a predicted secondary structure (Fig. 1a)^25^, which is also nearly identical to that of the experimentally determined structure of the AcrA from *E.coli,* with the exception of the divergent C-terminal region, which is predicted to be disordered and is not seen in the available cryo-EM structures^8, 9^. This allows creation of reliable homology models of AcrE, which we have previously reported^25^. In contrast to AcrA and AcrE, both MdtA and MdsA are more sequentially divergent, with MdsA sharing less than 30% identity with AcrA and AcrE, which is predicted to also translate in differences into significant differences 3D structure^5^.

**Figure 1.**
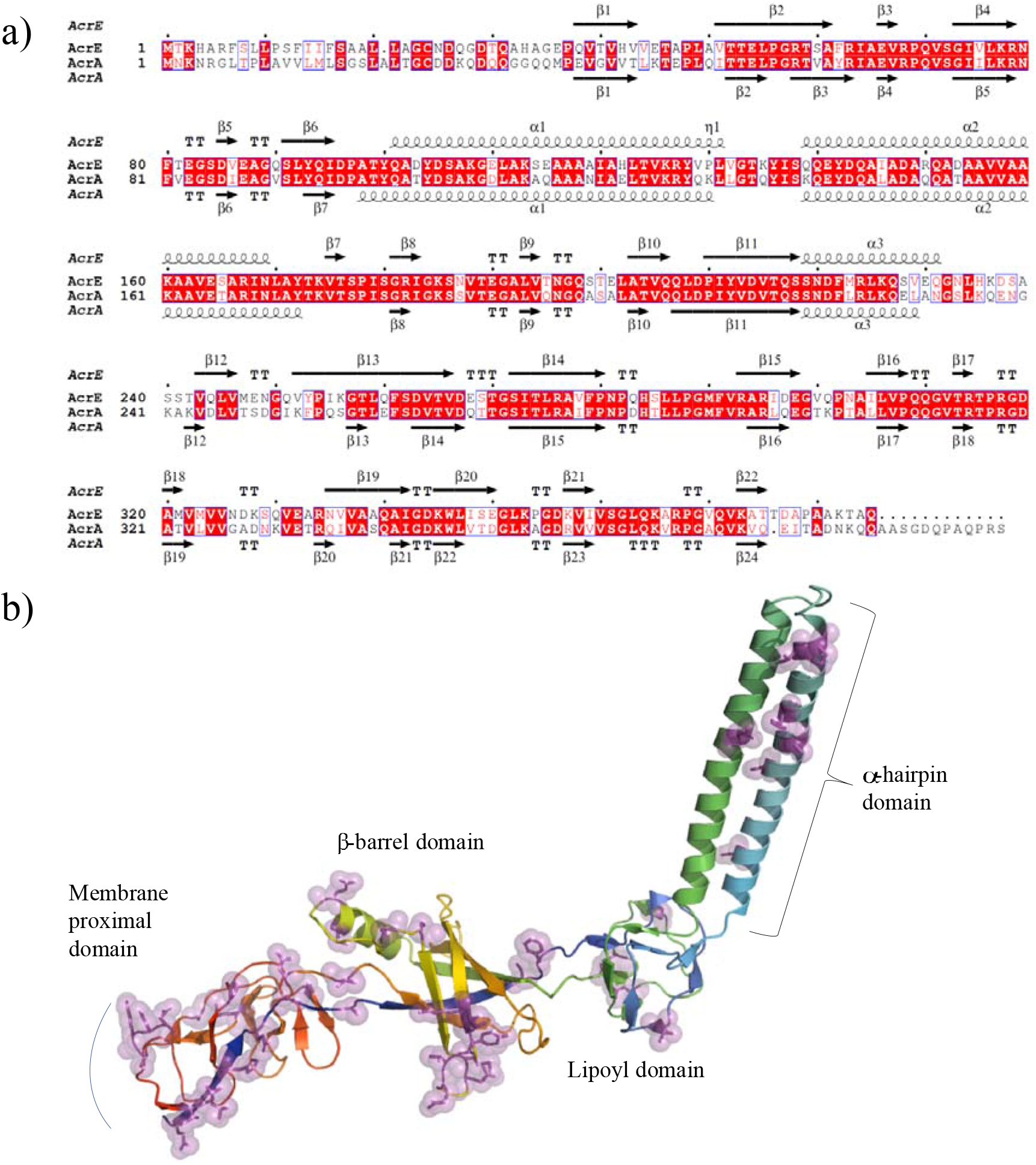
**a**) A pairwise sequence alignment of AcrA and AcrE of *S. enterica* highlighting their predicted close structural homology. The top secondary structure is derived from the previously reported homology model of AcrE^25^, while the bottom secondary structure corresponds to the experimental AcrA structure from *E. coli* (PDB ID 5O66; chain G), which has no sequence gaps with the AcrA of *S. enterica*. **b)** Mapping the sequence differences between the *Salmonella* AcrE and AcrA, onto the homology model of the AcrE^25^. The non-conserved substitutions are shown in sidechain and semi-transparent sphere representation. The mapping demonstrates that the bulk of the discrepancies, which may be expected to account for the functional differences between the PAPs map to their beta-barrel and membrane-proximal domains.

Although AcrA is the cognate PAP for AcrB, the RND pump AcrD was shown to depend on AcrA to form a functional tripartite efflux system since it lacks is not encoded with its own PAP^26^. Indeed, AcrA has been reported to also function with AcrF in *E. coli*^27^ and recently, AcrE has been demonstrated to function with AcrB in *S.* Typhimurium^25^.

The major RND-transporter binding residues of AcrA have been highlighted by cryogenic electron microscopy structural studies^8, 9^ and validated by mutagenesis^25^. Our comparative analysis of *Salmonella* PAPs demonstrated that these critical residues fall within a discrete number of linear sequence sites, which we termed RND-binding boxes^25^. These are shared between AcrA and AcrE, potentially explaining their interchangeability^25^. However, MdtA and MdsA are not interchangeable and cannot function with non-cognate RND-transporters and correspondingly are significantly different within RND-binding boxes^25^. Although AcrA and AcrE have been shown to be largely interchangeable, the ability of AcrE to function with AcrD remains unknown.

Here, we have investigated the substrate specificity of *S.* Typhimurium SL1344 AcrD. We furthermore explored whether the interoperability of AcrA and AcrE extends to the RND-transporter AcrD and whether this interaction is driven by the same residues that have been shown to be important for other PAP-RND combinations.

## Materials and methods

### Bacterial strains

All strains used in this study are listed in Table 1. The *Salmonella enterica* serovar Typhimurium strains were derived from the wild-type strain SL1344 (henceforth referred to as *S.* Typhimurium), a pathogenic strain first isolated from an experimentally infected calf^28^. All strains were grown in Luria–Bertani (LB) broth at 37°C with aeration.

**Table 1.**
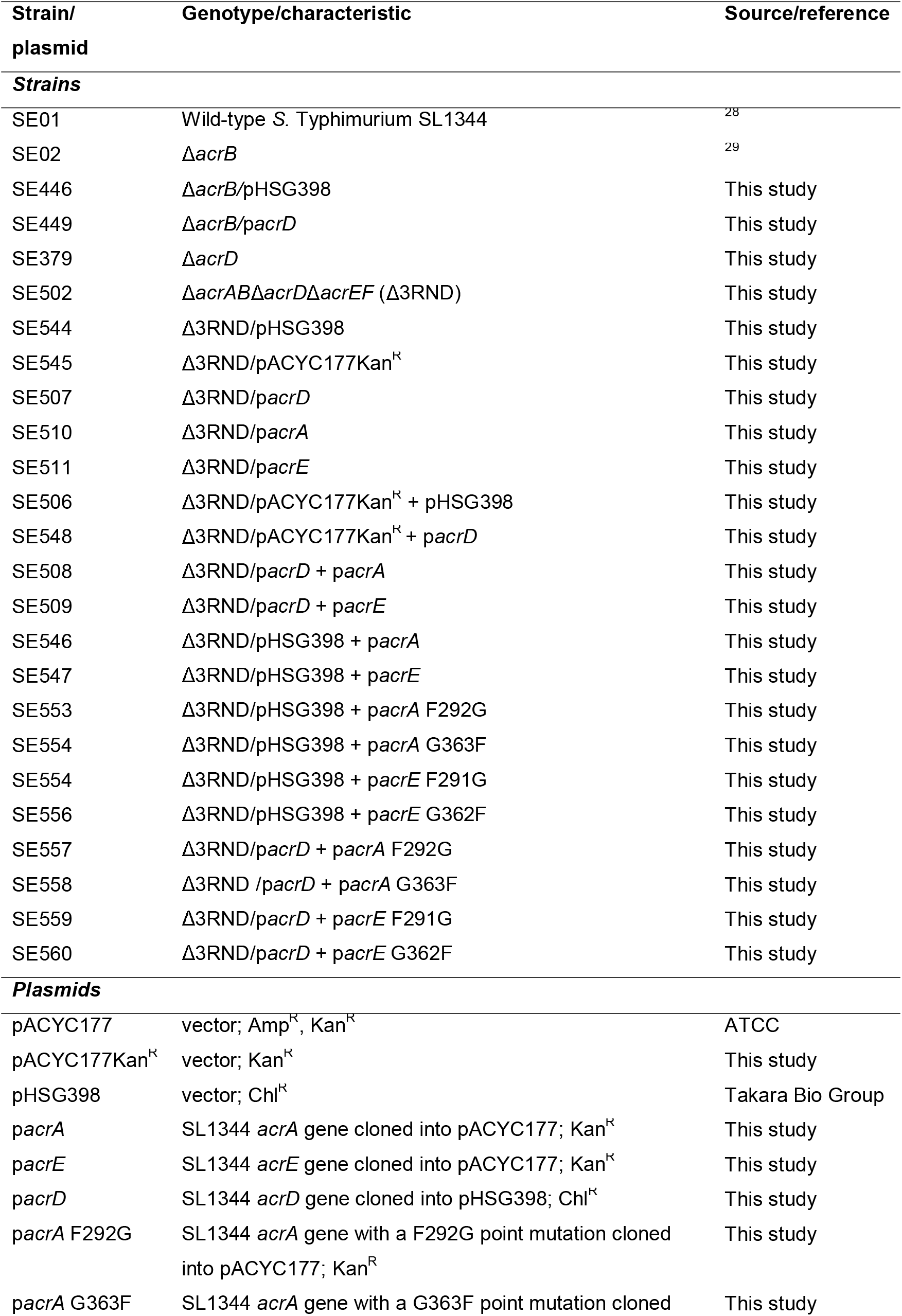
List of *S. enterica* serovar Typhimurium strains and plasmids used in this study.

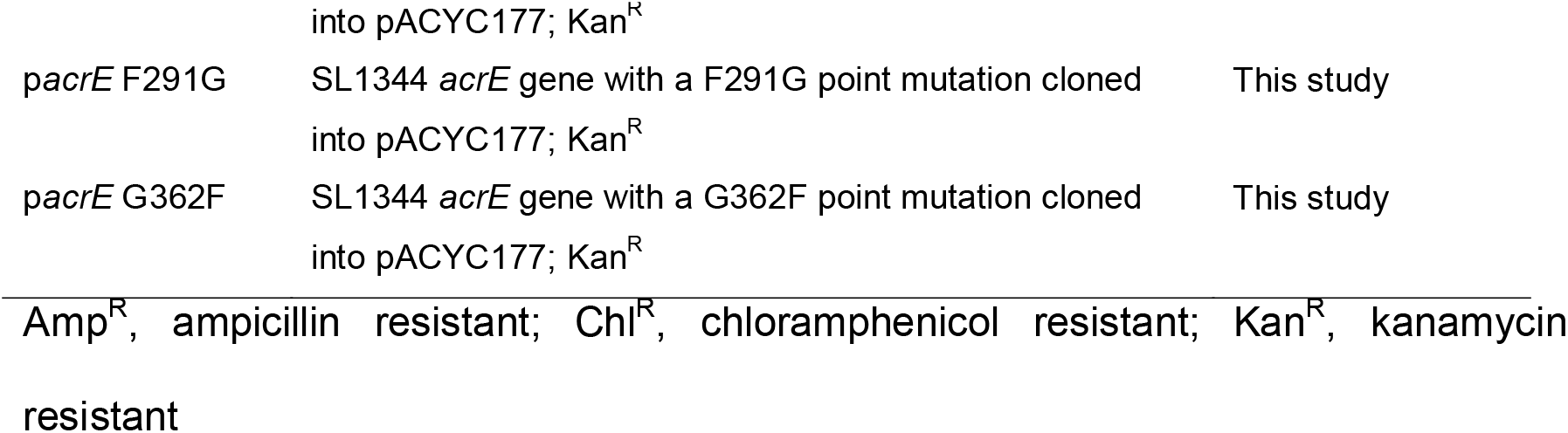

### Construction of gene deletion mutants

The Δ*acrB* mutant strain was constructed previously ^29^. All other mutant strains were constructed using the λ red recombinase system described previously, antibiotic markers were removed, and the process repeated to make double and triple knockout *S.* Typhimurium SL1344 strains (Table 1)^30^. All the primers used for generating gene knockouts and cloning are listed in Table S1.

### Plasmid construction

All plasmids used in this study are listed in Table 1. The *acrD* and *acrA* genes were amplified from *S.* Typhimurium SL1344 by PCR and cloned into pHSG398 and pACYC177 plasmids, respectively, as described previously^26^. Expression of the *acrE* gene is repressed by H-NS ^15^. Therefore, to clone *acrE* into pACYC177 and obtain sufficient expression, a forward primer was designed containing the *trc* promoter and the *acrE* ribosomal binding site (Table S1). The synthetic *trc* promoter is derived from the *E. coli trp* and *lac*UV5 promoters that drives high level of transcription^31^.

The *acrE* gene was amplified from *S.* Typhimuirum SL1344 genomic DNA by PCR using the *acrE* cloning F and R primers (Table S1), which introduced *Sca*I and *Bam*HI sites, respectively. The PCR fragment contained the *trc* promoter and a region 14 base pairs upstream to 2 base pairs downstream of *acrE*. This fragment was digested with *Sca*I and *Bam*HI and cloned into the corresponding sites of pACYC177, where an ampicillin resistance gene was located. The resulting plasmid pACYC177 *acrE* solely possessed a kanamycin resistance marker. The control pACYC177Kan^R^ plasmid was constructed as described previously^26^.

### Construction of mutant p*acrA* and p*acrE* plasmids

The *acrA* and *acrE* point mutants were generated using the GeneArt^⍰^ Gene Synthesis Service (Invitrogen, Germany) and subsequently cloned into the pACYC177 plasmid using the Subcloning Service (Invitrogen, Germany). All plasmids were sequenced to check for the presence of the desired point mutations and to ensure there were no unwanted secondary mutations.

### Determination of antimicrobial susceptibility

The minimum inhibitory concentration (MIC) of various antimicrobials was determined using the agar dilution method according to CLSI guidance^32^.

## Results and discussion

### AcrD of *S.* Typhimurium SL1344 does not transport aminoglycosides

Despite being isolated several decades ago^33^, the substrate specificity of AcrD remains relatively poorly characterised experimentally. Therefore, an additional rationale of this study was to investigate the substrate specificity of *S.* Typhimurium SL1344 AcrD, especially in the context of PAP-RND interactions, which may provide modulatory effects on the specificity of the pump. Previously, it has been reported that *E. coli* AcrD exports aminoglycosides^19, 20^. However, there is a lack of experimental evidence in *Salmonella* and most of the features of *Salmonella* AcrD are inferred based on close sequence similarity to *E. coli* AcrD (97.4%, Fig. S1). While, some previous work has addressed this, aminoglycosides have not been specifically investigated^5, 17^. Therefore, we investigated the substrate range of AcrD in *S.* Typhimurium SL1344.

The Δ*acrD* SL1344 strain did not exhibit any significant increase in susceptibility to any of the antimicrobials tested as previously reported^17^. This is likely because expression of *acrD* is generally low in laboratory conditions and for many compounds, any effect would be masked by the presence of AcrB^17^. Therefore, p*acrD* was transformed into the Δ*acrB* strain, and the effect of *acrD* overexpression on antimicrobial susceptibility of the resulting transformant was determined. The Δ*acrB*/p*acrD* strain displayed significantly increased MIC values to reported AcrD-substrates aztreonam, carbenicillin, cloxacillin, fusidic acid, nafcillin, novobiocin and oxacillin (Table 2) consistent with previous studies^26^, suggesting that protein is functionally expressed and incorporated into the membrane. Surprisingly, the introduction of *acrD* into Δ*acrB (*Δ*acrB*/p*acrD)* strain did not result in a significant increase in MICs to the aminoglycosides kanamycin, gentamicin, spectinomycin or streptomycin (Table 2), implying that AcrD is not measurably contributing to aminoglycoside efflux. This is in contrast to the reported role of AcrD in the aminoglycoside resistance of *E. coli*, wherein deletion of *acrD* was shown to decrease aminoglycoside MICs by two to eight fold^19^ and expression of *acrD* from a plasmid in an *acrB::aph* Δ*acrD* strain increased aminoglycoside MICs by two fold^34^. In agreement with our findings, the AcrD efflux pump of the Gram-negative plant pathogen *Erwinia amylovora* has also been reported to not play a role in aminoglycoside resistance^35^. Similarly, our data shows that the AcrD efflux pump of *S.* Typhimurium SL1344 does not seem to be involved in aminoglycoside export as previously assumed.

**Table 2.** Susceptibility of *S.* Typhimurium strains to antimicrobials.

| Strain | MIC (mg/L) |  |  |  |  |  |  |  |  |  |  |  |
| --- | --- | --- | --- | --- | --- | --- | --- | --- | --- | --- | --- | --- |
|  | ATM | CAR | CXA | FA | NAF | NOV | OXA | TIC | GEN | SPR | STR | KAN |
| Wild type SL1344 | 0.06 | 4 | 512 | 1024 | 1024 | 512 | 512 | 4 | 0.5 | 16 | 8 | 1 |
| Δ*acrB* | 0.06 | 1 | 4 | 4 | 8 | 2 | 4 | 1 | 0.25 | 16 | 8 | 1 |
| Δ*acrB*/pHSG398 | 0.06 | 1 | 4 | 4 | 8 | 2 | 2 | 1 | 0.25 | 16 | 4 | 1 |
| Δ*acrB*/*pacrD* | **0.25** | **8** | **16** | **64** | **64** | **8** | **16** | **16** | 0.25 | 16 | 4 | 1 |
| Δ*acrD* | 0.06 | 4 | 512 | 1024 | 1024 | 512 | 512 | 4 | 0.5 | 16 | 4 | 1 |
| Δ*acrAB*Δ*acrD*Δ*acrEF* (Δ3RND) | 0.06 | 1 | 1 | 4 | 2 | 1 | 1 | 1 | 0.25 | 16 | 4 | 1 |
| Δ3RND/pHSG398 | 0.06 | 0.5 | 1 | 4 | 2 | 1 | 1 | 1 | 0.25 | 16 | 4 | 1 |
| Δ3RND/pACYC177Kan<sup>R</sup> | 0.06 | 1 | 1 | 4 | 2 | 1 | 1 | 1 | 0.5 | 16 | 4 | >32 |
| Δ3RND/*pacrD* | 0.06 | 0.5 | 1 | 4 | 2 | 1 | 1 | 1 | 0.25 | 16 | 4 | 1 |
| Δ3RND/*pacrA* | 0.06 | 1 | 1 | 4 | 2 | 1 | 1 | 1 | 0.5 | 16 | 4 | >32 |
| Δ3RND/*pacrE* | 0.06 | 1 | 1 | 4 | 2 | 1 | 1 | 1 | 0.5 | 16 | 4 | >32 |
| Δ3RND/pACYC177Kan<sup>R</sup> + pHSG398 | 0.06 | 0.5 | 1 | 4 | 2 | 1 | 1 | 1 | 0.25 | 16 | 8 | >32 |
| Δ3RND/pACYC177Kan<sup>R</sup> + *pacrD* | 0.06 | 1 | 1 | 4 | 2 | 1 | 1 | 1 | 0.25 | 16 | 4 | >32 |
| Δ3RND/*pacrD* + *pacrA* | **0.25** | **8** | **16** | **128** | **32** | **8** | **16** | **8** | 0.5 | 16 | 4 | >32 |
| Δ3RND/*pacrD* + *pacrE* | **0.25** | **8** | **16** | **128** | **32** | **8** | **16** | **8** | 0.5 | 16 | 4 | >32 |
| Δ3RND/pHSG398 + *pacrA* | 0.06 | 0.5 | 1 | 4 | 2 | 1 | 1 | 1 | 0.5 | 16 | 4 | >32 |
| Δ3RND/pHSG398 + *pacrE* | 0.06 | 0.5 | 1 | 4 | 2 | 1 | 1 | 1 | 0.5 | 16 | 4 | >32 |
| Δ3RND/pHSG398 + F292G *acrA* | 0.06 | 0.5 | 1 | 4 | 2 | 1 | 1 | 1 | 0.5 | 16 | 4 | >32 |
| Δ3RND/pHSG398 + G363F *acrA* | 0.06 | 0.5 | 1 | 4 | 2 | 1 | 1 | 1 | 0.5 | 16 | 4 | >32 |
| Δ3RND/pHSG398 + F291G *acrE* | 0.06 | 0.5 | 1 | 4 | 2 | 1 | 1 | 1 | 0.5 | 16 | 4 | >32 |
| Δ3RND/pHSG398 + G362F *acrE* | 0.06 | 0.5 | 1 | 4 | 2 | 1 | 1 | 1 | 0.5 | 16 | 4 | >32 |
| Δ3RND/*pacrD* + F292G *acrA* | 0.06 | 0.5 | 1 | 4 | 2 | 1 | 1 | 1 | 0.5 | 16 | 4 | >32 |
| Δ3RND/*pacrD* + G363F *acrA* | 0.06 | 1 | 1 | 4 | 2 | 1 | 1 | 1 | 0.25 | 16 | 4 | >32 |
| Δ3RND/*pacrD* + F291G *acrE* | 0.06 | 0.5 | 1 | 4 | 2 | 1 | 1 | 1 | 0.25 | 16 | 4 | >32 |
| Δ3RND/*pacrD* + G362F *acrE* | 0.06 | 1 | 1 | 4 | 2 | 1 | 1 | 1 | 0.25 | 16 | 4 | >32 |

ATM, aztreonam; CAR, carbenicillin; CXA, cloxacillin; FA, fusidic acid; GEN, gentamicin;
KAN, kanamycin; NAF, nafcillin; NOV, novobiocin; SPR, spectinomycin, STR, streptomycin, TIC,
ticarcillin
Values in bold indicate a significant increase (>2-fold) than those of their corresponding parental
strains.

A possible explanation for the differences in the substrate profiles of AcrD between *E. coli* and *S.* Typhimurium could be the observed discrepancy between the residues in their respective access and deep binding pockets (Fig. S1). Due to the lack of experimental AcrD structure, the functional significance of the residues of the respective drug binding pockets of AcrD is inferred from their positional homology with corresponding AcrB residues, structures of which have been experimentally defined for both *E. coli* ^36–38^, and more recently for *Salmonella*^39^ Specifically, the presence of a serine in the deep binding pocket of *S.* Typhimurium AcrD at position 610, which in *E. coli* AcrD is occupied by an alanine, could possibly impact the previously described lipophilic character of the drug binding cavity^22^. There are also two additional discrepancies which could be seen as non-conservative substitutions, namely that of *E. coli* AcrD isoleucine to a phenylalanine at position 633 (I633F) in *S.* Typhimurium, and leucine to a glutamine at position 565 (L565Q), both of which are likely to cause steric hinderance and impact the electrostatics of the access binding pocket, respectively^21, 22^. These subtle differences may account for the notable differences in substrate recognition by AcrD between the two species.

### AcrE forms a functional PAP-RND pair with AcrD

AcrD has been previously shown to depend on AcrA to function as an efflux system^26^. Therefore, owing to the high similarity of the predicted RND-binding sites between the PAPs AcrA and AcrE^25, 40^, we hypothesised that AcrE should also function with AcrD. To test this, we deleted the *acrAB*, *acrD* and *acrEF* genes in *S.* Typhimurium SL1344 to give a strain without active RND-dependent efflux, as indicated by significantly increased susceptibility to AcrB-, AcrF-, and AcrD- substrates (Table 2 and S2). The MdtABC and MdsABC systems are much less similar to the three AcrB/AcrD/AcrF-based systems and play a minor role in resistance. Consistent with this, they are not expressed under standard laboratory conditions^5^ and furthermore their inactivation did not have any additive effect on antimicrobial susceptibility^5, 25^. Hence, these systems were not inactivated.

Firstly, we validated the previously reported AcrA dependency of AcrD in *S.* Typhimurium SL1344^26^. The p*acrA* and p*acrD* plasmids were co-transformed into the Δ3RND strain, and the antimicrobial susceptibility of the resulting transformant was determined. We found that co-expression of *acrA* and *acrD* in the Δ3RND strain significantly decreased susceptibility to known AcrD-substrates aztreonam, carbenicillin, cloxacillin, fusidic acid, nafcillin, novobiocin, oxacillin and ticarcillin (Table 2).

Secondly, to determine whether AcrE and AcrD form a functional complex together, p*acrD* and p*acrE* were co-transformed into the Δ3RND strain and the susceptibility to validated AcrD-substrates was tested. Co-expression of *acrE* and *acrD* in the Δ*3RND* strain significantly increased the MICs of aztreonam, carbenicillin, cloxacillin, fusidic acid, nafcillin, novobiocin, oxacillin and ticarcillin (Table 2). There was no difference in MIC values between co-expressing *acrD* with *acrA* or *acrE*, which demonstrates the full interchangeability of the two PAPs (Table 2). Furthermore, co-expression of either *acrE* and *acrD* or *acrA* and *acrD* in the Δ3RND strain did not increase MIC values to the tested AcrB-substrates (i.e., acriflavine, crystal violet, ethidium bromide, erythromycin, methylene blue, rhodamine 6G and tetracycline)^5^, clearly showing AcrD-mediated efflux (Table S2). Overexpression of either *acrD* or *acrE* alone in the Δ3RND strain did not significantly increased MIC values to the AcrD-substrates tested (Table 2), signifying that AcrE requires the presence of AcrD to form a functional, efflux-competent complex.

Our data suggests interchangeability between AcrA and AcrE in *S.* Typhimurium SL1344. One possible explanation for the interoperability between AcrA and AcrE is that the latter may function as a backup PAP for when AcrA function is impaired or lost. This idea is supported by evidence from studies that demonstrated that in *S.*

Typhimurium, in the absence of *acrA*, it was possible to select for *acrE* overexpression^25 41^. Another study demonstrated that in the absence of *acrA* and *acrE*, it is possible to restore the phenotypic defect in active efflux by complementing with either *acrA* or *acrE*^40^.

### Disruption of the RND-binding residues in AcrA or AcrE impairs AcrD-mediated efflux of substrate drugs

Previously, we showed that the promiscuity between *Salmonella* AcrA and AcrE in their ability to form a functional complex with AcrB stems from the highly conserved RND-binding sites (termed RND-binding boxes) between these two PAPs. Specifically, within the *Salmonella* AcrA, we identified several residues mapping to the β-barrel and membrane proximal domains that were important for AcrB-binding^25^. There, the disruption of the F292 or G363 residues in AcrA produced the most pronounced phenotypic effect, resulting in severely abrogated active efflux and significantly increased susceptibility to AcrB-substrates^25^. Therefore, to investigate whether these residues are also important for binding of the newly determined cognate PAPs to AcrD, the point mutation corresponding to F292G or G363F were constructed in both p*acrA* and p*acrE* (F291G and G362F) respectively and co-transformed with p*acrD* into the Δ3RND strain. Based on structural analysis, we chose F292G and G363F as target mutations due to their radical change of respective site-chain properties.

Consistent with the data obtained in coexpression with AcrB^25^, the disruption of F292 or G363 in AcrA resulted in impaired AcrD-mediated efflux of AcrD-substrates, confirming that the same residues required for binding of AcrA to AcrB are also required for its binding to AcrD (Table 2). These point mutations do not impact the protein levels and folding as previously demonstrated^25^. To determine whether the corresponding residues in AcrE are also important for AcrD-binding, F291 and G362 were mutated (Fig. S2). As expected, the F291G or G362F point mutations in AcrE also impaired AcrD-mediated efflux in the Δ3RND strain (Table 2). These data suggest that the PAP-RND binding sites previously identified based upon AcrA-AcrB interaction, are indeed both sequentially and functionally conserved between AcrA and AcrE and account for the productive recognition and formation of functional tripartite pumps.

### Concluding remarks

Here, we report that the PAP AcrE can form a functional complex with the RND-transporter AcrD, further validating the interchangeability between the homologous PAPs AcrA and AcrE. Furthermore, this interchangeability is likely to be due to the highly conserved and specific RND-binding sites between these two PAPs. Our report highlights the redundancy between these two PAPs must be taken into account when targeting them for efflux inhibition. Therefore, the residues we identified here could inform future design of effective efflux inhibitors targeting PAPs or tripartite complex assemblies.

## Transparency declaration

None to declare.

