## Supplementary data for "Interchangeability of Periplasmic Adaptor Proteins AcrA and AcrE in forming functional efflux pumps with AcrD in *Salmonella* Typhimurium"

**Table S1.** List of primers used in this study.

| **Primer description** | **Primer sequence (5’-3’)** |
| --- | --- |
| *acrAB* knockout forward | GTCACACTAAAAACGGAACCACTGCAAATCACAACTGAACGTGTAGGCTGGAGCTGCTTC |
| *acrAB* knockout reverse | GCTTGCGCGGCCTTATCAACAGTGAGCAAATCAGCGATGTGGGAATTAGCCATGGTCCAT |
| *acrD* knockout forward | GGTGCTGGCTATCCTGTTGTGTCTGACAGGGGCGTTGTGTAGGCTGGAGCTGCTTC |
| *acrD* knockout reverse | AAGCGGCGACGTATCAGCACGAAAAACAGGGGTACAGGGAATTAGCCATGGTCCAT |
| *acrE* knockout forward | GGTTTTCACTCCTGCCCTCATTCATCATATTCTCTGCTGCGTGTAGGCTGGAGCTGCTTC |
| *acrE* knockout reverse | CTTCGACCTGGCTTTTATCGTTAACCACCATCACCATTGCGGGAATTAGCCATGGTCCAT |
| *acrF* knockout forward | ACTACCGATGCTCCTGCAGCGAAAACGGCGCAATAAGGTAACCGTACATGGTGTAGGCTGGAGCTGCTTC |
| *acrF* knockout reverse | AAGGCGTCCGAAGACGCCTCTGTTTACCGGTTAATCATGATGGCGATTAAATGGGAATTAGCCATGGTCC |
| *acrD* cloning forward | CGCGGATCCCATTATCTCCTTTATTTCTCC |
| *acrD* cloning reverse | CGCGCATGCTTATTTCGGGCGCGGCTTCAG |
| *acrA* cloning forward | CGCAGTACTATGTCGGTGAATTTACAGGCG |
| *acrA* cloning reverse | CGCGGATCCGTCTTAACGGCTCCTGTTTAA |
| *acrE* cloning forward | GCGCAGTACTGAGCTGTTGACAATTAATCATCCGGCTCGTATAATGTGTGGAATTTTAAGGAACAGTAATGACG |
| *acrE cloning* reverse | GCGCGGATCCCCTTATTGCGCCGTTTTCGC |

The introduced restriction sites are underlined.

**Table S2.** Susceptibility of *S.* Typhimurium strains to known AcrB-substrates.

|  | **MIC (mg/L)** | | | | | | | |
| --- | --- | --- | --- | --- | --- | --- | --- | --- |
| **Strain** | **EB** | **ERY** | **CV** | **R6G** | **ACR** | **MB** | **CTX** | **TET** |
| Wild type SL1344 | >1024 | 128 | 64 | 1024 | 128 | 1024 | 0.12 | 1 |
| Δ*acrB* | 16 | 2 | 2 | 8 | 16 | 4 | 0.02 | 0.25 |
| Δ*acrB/*pHSG398 | 16 | 1 | 2 | 8 | 8 | 2 | 0.02 | 0.25 |
| Δ*acrB/*p*acrD* | 16 | 1 | 2 | 8 | 8 | 4 | 0.02 | 0.25 |
| Δ*acrD* | >1024 | 128 | 64 | 1024 | 128 | 1024 | 0.12 | 1 |
| Δ*acrAB*Δ*acrD*Δ*acrEF* (Δ3RND) | 8 | 2 | 2 | 4 | 8 | 4 | 0.02 | 0.25 |
| Δ3RND/pHSG398 | 16 | 1 | 4 | 4 | 8 | 2 | 0.02 | 0.25 |
| Δ3RND/pACYC177Kan^R^ | 16 | 2 | 2 | 4 | 8 | 2 | 0.02 | 0.25 |
| Δ3RND/p*acrD* | 16 | 2 | 2 | 4 | 8 | 4 | 0.02 | 0.25 |
| Δ3RND/p*acrA* | 16 | 2 | 2 | 4 | 8 | 4 | 0.02 | 0.25 |
| Δ3RND/p*acrE* | 16 | 2 | 2 | 4 | 8 | 2 | 0.02 | 0.25 |
| Δ3RND/pACYC177Kan^R^ + pHSG398 | 16 | 2 | 2 | 4 | 8 | 2 | 0.02 | 0.25 |
| Δ3RND/pACYC177Kan^R^ + p*acrD* | 8 | 1 | 2 | 4 | 4 | 2 | 0.02 | 0.25 |
| Δ3RND/p*acrD* + p*acrA* | 16 | 2 | 2 | 4 | 8 | 4 | 0.02 | 0.25 |
| Δ3RND/p*acrD* + p*acrE* | 16 | 2 | 2 | 4 | 8 | 4 | 0.02 | 0.25 |
| Δ3RND/pHSG398 + p*acrA* | 16 | 1 | 4 | 4 | 8 | 2 | 0.02 | 0.25 |
| Δ3RND/pHSG398 + p*acrE* | 16 | 1 | 2 | 4 | 8 | 2 | 0.02 | 0.25 |
| Δ3RND/pHSG398 + F292G *acrA* | 16 | 1 | 2 | 4 | 8 | 2 | 0.02 | 0.25 |
| Δ3RND/pHSG398 + G363F *acrA* | 16 | 1 | 2 | 4 | 8 | 2 | 0.02 | 0.25 |
| Δ3RND/pHSG398 + F291G *acrE* | 16 | 1 | 2 | 4 | 8 | 2 | 0.02 | 0.25 |
| Δ3RND/pHSG398 + G362F *acrE* | 16 | 1 | 2 | 4 | 8 | 2 | 0.02 | 0.25 |
| Δ3RND/p*acrD* + F292G *acrA* | 16 | 1 | 2 | 4 | 8 | 2 | 0.02 | 0.25 |
| Δ3RND /p*acrD* + G363F *acrA* | 16 | 1 | 2 | 4 | 8 | 2 | 0.02 | 0.25 |
| Δ3RND/p*acrD* + F291G *acrE* | 16 | 1 | 2 | 4 | 8 | 2 | 0.02 | 0.25 |
| Δ3RND/p*acrD* + G362F *acrE* | 16 | 1 | 2 | 4 | 8 | 2 | 0.06 | 0.25 |

ACR, acriflavine; CTX, cefotaxime; CV, crystal violet; EB, ethidium bromide; ERY, erythromycin; MB, methylene blue; R6G, rhodamine 6G; TET, tetracycline

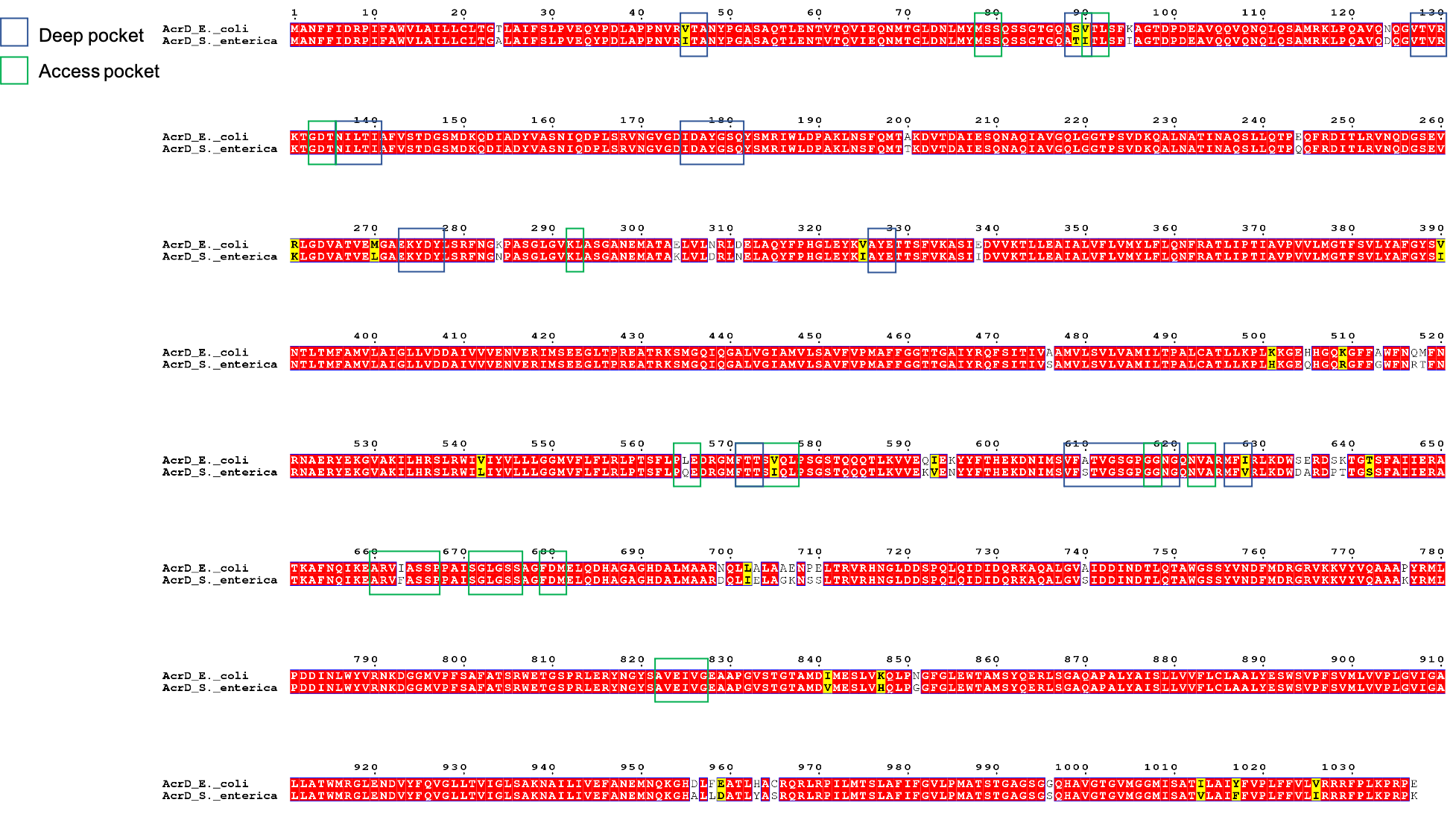

**Figure S1. Pairwise sequence alignment of AcrD of *E. coli* K-12 and AcrD of *S.* Typhimurium SL1344.** Identical residues are highlighted in red, similar in yellow, while all others are mismatches. Green and blue boxes indicate residues in the access pocket and deep pocket, respectively^1^.

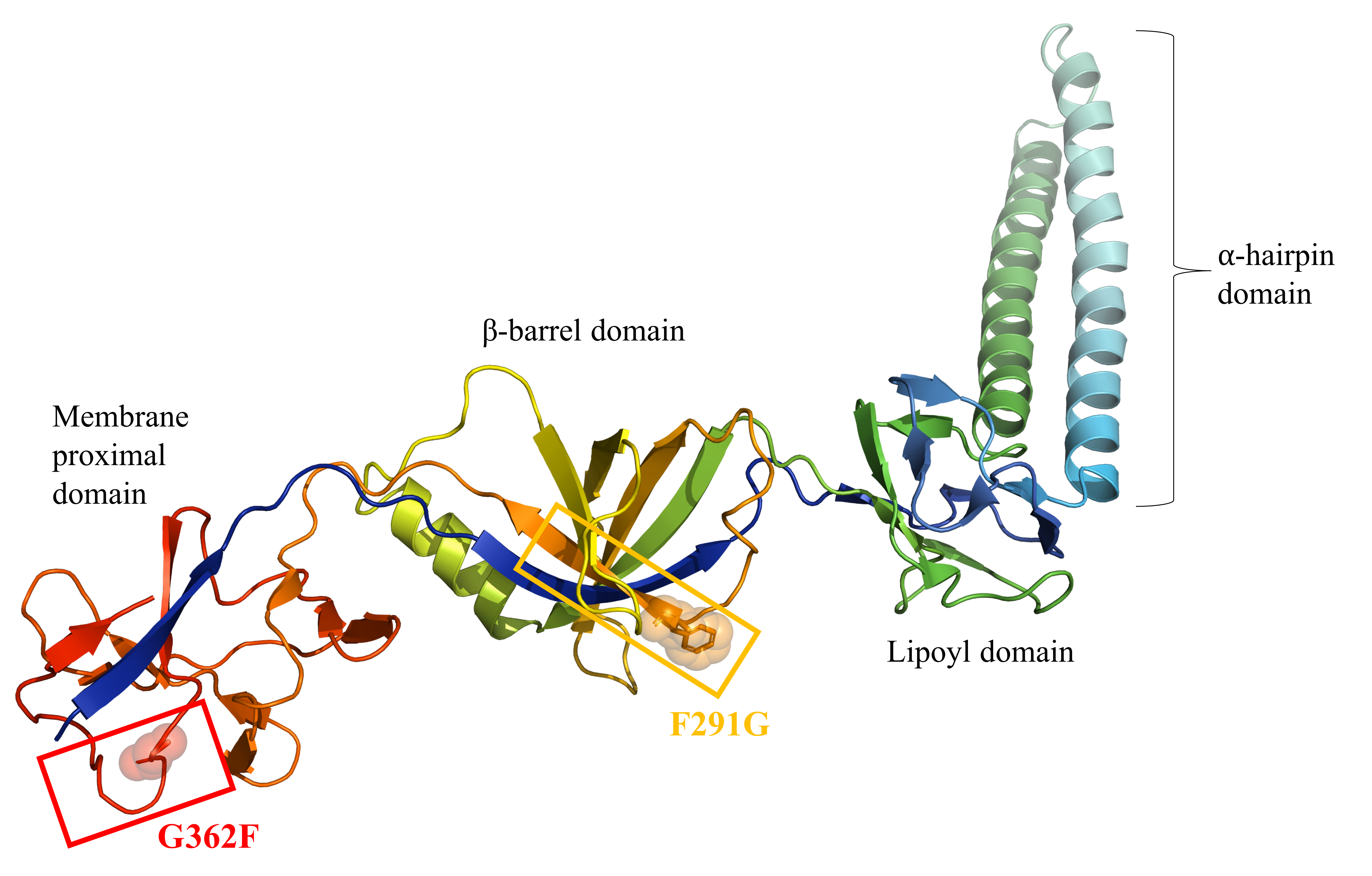

**Figure S2. A homology model of the structure of AcrE showing the positions of residues F291 and G362 located in the RND-binding boxes 5 and 9 respectively^2^, which are shown to be responsible for AcrD-interaction by our site-directed mutagenesis.** The position of the binding-box 5 is shown in orange, while box 9 is delineated by a red box. It is notable that despite the large discrepancies in the primary sequences of AcrA and AcrE (as seen in Fig.1a), the loops and β-strands that form these boxes are significantly conserved between AcrA and AcrE.
